# *In vivo*-mimicking 3D cultures secrete distinct extracellular vesicles upon cancer cell invasion

**DOI:** 10.1101/2020.12.22.423913

**Authors:** Jens C. Luoto, Leila S. Coelho-Rato, Sara H. Bengs, Jannica Roininen, John E. Eriksson, Lea Sistonen, Eva Henriksson

## Abstract

Extracellular vesicles (EVs) loaded with biomolecules are important in intercellular communication and mediate local and long-range signals in cancer metastasis. However, it is currently unknown how the development of the primary tumor and onset of invasion affect the secretion and characteristics of EVs. In this study, we developed an EV production method utilizing *in vivo*-mimicking extracellular matrix-based 3D cultures, which allows tracking of EVs over the course of invasive development of tumor organoids. Using this method, combined with proteomic profiling, we show that PC3 human prostate cancer organoids secrete EVs with previously undefined protein cargo, which substantially differs from EV cargo of 2D cultured cells. Intriguingly, an increase in EV amounts and extensive changes in EV protein composition were detected upon invasive transition of the organoids. These results reveal that EV secretion and cargo loading are highly dependent on the developmental status of the tumor organoid, emphasizing the necessity of *in vivo*-mimicking conditions for discovery of novel cancer-derived EV components, applicable as diagnostic markers for cancer.

## Introduction

The leading cause of cancer-related deaths worldwide is metastasis. In metastasis, cancer cells break free from the primary tumor and invade adjacent tissues. This is followed by intravasation into the bloodstream and extravasation into new tissues, forming secondary tumors (Hanahan and Weinberg, 2011). It is crucial to detect tumors before the onset of invasion, to prevent the metastasis cascade and reduce cancer mortality. Tumor progression and metastasis depend strongly on intercellular communication, in which extracellular vesicles (EVs) play a key role by shuttling biomolecules, such as proteins, lipids and nucleic acids across cells and tissues (Yáñez-Mó *et al*., 2015). Cells secrete EVs of different sizes, such as exosomes (30-100 nm), microvesicles (50-1000 nm), and larger cancer-specific oncosomes (up to 10 μm) (Van Niel, D’Angelo and Raposo, 2018). Cancer-derived EVs have been shown to promote metastasis by increasing angiogenesis, suppressing the immune system, reprogramming fibroblasts, and preparing pre-metastatic niches in specific organs (Kogure, Yoshioka and Ochiya, 2020). Cancer-derived EVs have evoked great interest as biomarkers for early detection of cancer, since they bear distinct cargo components and they are detected in most human biofluids, including blood, milk, and urine (Yáñez-Mó *et al*., 2015; Yamamoto, Kosaka and Ochiya, 2019; Zhang and Yu, 2019; Pang *et al*., 2020). However, biofluids contain heterogeneous and complex populations of EVs secreted by different types of cells, and patient samples are often derived from late-stage cancer patients. Therefore, novel methods simulating tumor growth in a more controlled context would be acutely needed for cancer EV research.

Previous studies on cancer EVs have extensively used conventional two-dimensional (2D) cell cultures (Xu *et al*., 2018), in which *in vivo* tumor growth and development are poorly recapitulated (Duval *et al*., 2019). Due to the lack of microenvironment, which normally exists in tissues and organs, cell-cell and cell-extracellular matrix (ECM) contacts are aberrant in 2D culture conditions. In contrast, three-dimensional (3D) cell cultures allow cancer cells to acquire similar morphology, growth and differentiation patterns as detected in tumors. There are currently multiple 3D culture approaches to grow organoids, in contrast to the standardized and comparable 2D culture systems. Specifically, secretion and cargo components of EVs differed substantially between 2D and 3D cultures (Eguchi *et al*., 2018; Rocha *et al*., 2019; Thippabhotla, Zhong and He, 2019). However, these studies used non-modifiable 3D matrices, where the cancer cells are impeded from invading their surroundings, thus failing to mimic invasion. In case modifiable matrices were used in 3D, the analyses were limited to a single time point (Szvicsek *et al*., 2019; Yang, Knight and Stephens, 2020). Hence, it is unknown whether EV secretion and characteristics change during tumor development and invasive transition. It is of utmost importance to the EV field to find model systems that can emulate cancer development and invasion, enabling more efficient EV cancer biomarker discovery.

In this study, we developed an ECM-based 3D culture method for studying secretion of EVs from prostate cancer organoids over the course of their invasive development. Using this method and proteomic profiling, we uncovered previously undefined protein cargo and profound differences in cargo composition between EVs derived from 2D and 3D cultures. Most importantly, we found an increase in EV amounts and extensive changes in EV protein composition upon the invasive transition of the organoids. Our study demonstrates that EV secretion of cancer organoids in 3D *in vivo*-mimicking conditions is a dynamic process that changes according to the invasive state of the cells, which should not be neglected when investigating EVs in cancer biology.

## Results and Discussion

### *In vivo*-mimicking ECM-based 3D cultures enable EV isolation during invasive transition of tumor organoids

To address if the invasive development of tumors affects the secretion or characteristics of EVs, we established an *in vivo*-mimicking EV production method using organoid cultures undergoing invasive transition (Fig 1). We seeded human PC3 prostate cancer cells between two layers of the ECM hydrogel Matrigel and allowed the cells to divide and form organoids. The organoid-derived EVs were collected from the conditioned media every other day and the EVs were isolated by differential centrifugation into 10K and 100K pellets. The organoids underwent dynamic phenotypic changes during the culture period of 12 days. A major phenotypic change was detected from day 10 onwards, as cells invaded the matrix surrounding the organoids, forming clearly visible invasive structures (Fig 1). The timing of the invasive transition is in agreement with previous studies conducted with PC3 cells (Härmä *et al*., 2010; Björk *et al*., 2016).

**Figure 1.**
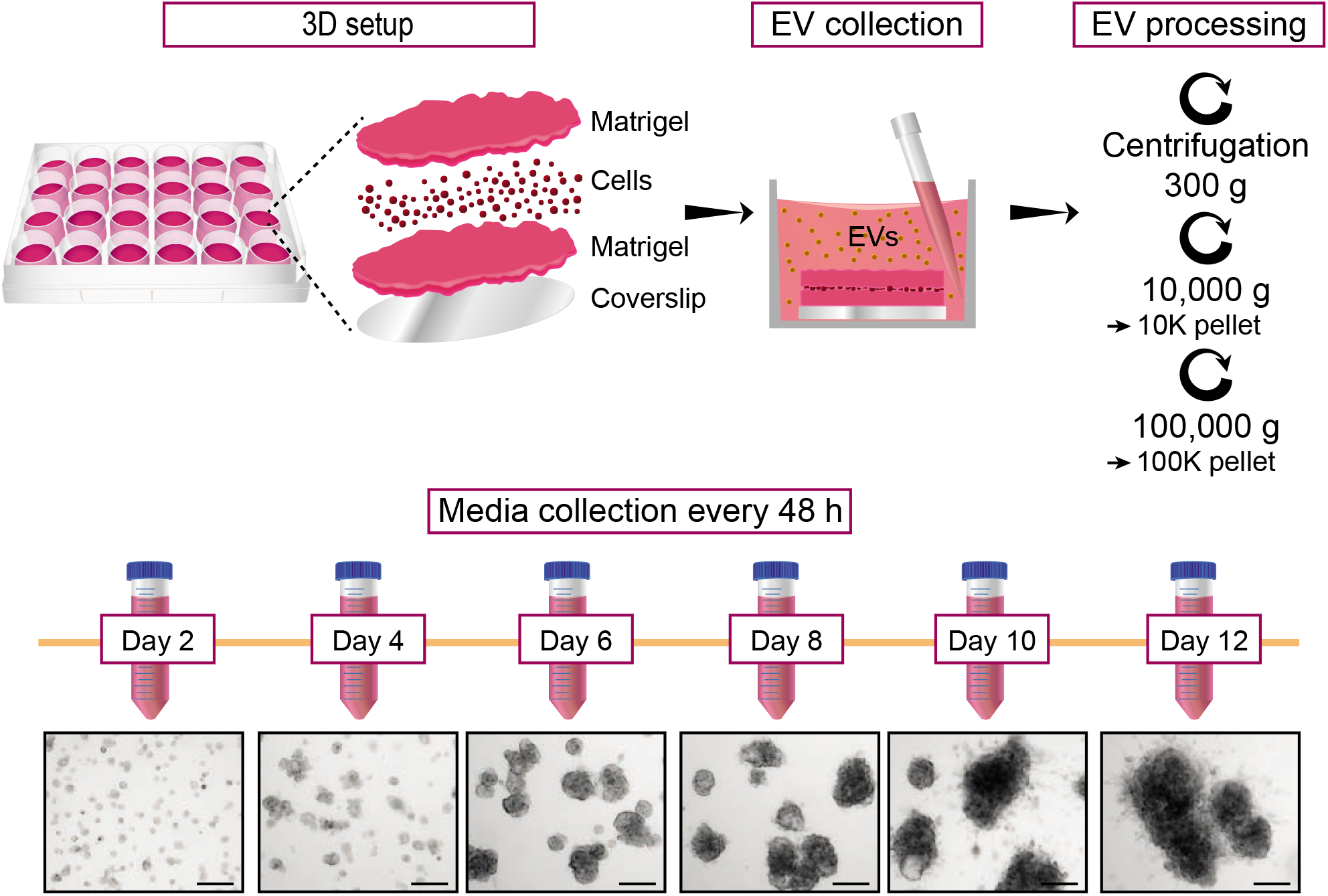
Isolation of EVs from ECM-based 3D cultures undergoing invasive transition. Graphical workflow of EV isolation from the ECM-based 3D cell culture. PC3 cells were seeded on top of coverslips in-between two layers of Matrigel (3.5 mg/ml) and allowed to grow and form organoids during 12 days. Conditioned media of the organoid cultures was collected every two days, from which EVs were isolated by differential centrifugation into 10K and 100K pellets. The organoids developed invasive structures from day 10 onwards, as shown by the representative brightfield images. Scale bars 200 μm.

Optimal conditions for EV isolation and PC3 organoid development were tested with three different Matrigel concentrations (3.5, 5.0, and 6.5 mg/ml) (Supplementary Fig 1A). In all cases, PC3 cells invaded the surrounding ECM and the cultures produced particles with similar sizes and amounts, as determined by nanoparticle tracking analysis (NTA) (Supplementary Fig 1B-D). The highest quantity of secreted particles was found in 3.5 mg/ml of Matrigel (Supplementary Fig 1C), and this concentration was chosen for all assays.

### The invasive transition of prostate cancer organoids causes a surge in EV secretion

To characterize the EVs secreted from the developing PC3 organoids undergoing invasive transition, we collected and isolated EVs into 10K and 100K pellets and conducted NTA and immunoblotting analyses (Fig 2A-C). Surprisingly, the 100K pellet samples, but not 10K, showed a significant increase in particle concentration at day 12 (Fig 2A), whereas the sizes of the particles were similar throughout the time course in both 10K and 100K samples (Fig 2B and Supplementary Fig 2). Similarly, immunoblotting showed markedly increased levels of selected EV markers; CD81 antigen (CD81), CD63 antigen (CD63) and tumor susceptibility gene 101 protein (TSG101) at day 10 onwards (Fig 2C). No major contamination was detected, as neither lipoproteins, e.g. apolipoprotein A1 (APOA1), nor intracellular proteins, e.g. lamin A/C and Golgi matrix protein (GM130), were present in the EV samples. In order to address if the sudden rise in the EV secretion was due to an abruptly increased number of cells, we counted the cells of the developing organoid cultures. Cells displayed a linear proliferation throughout the culture period of 12 days (Fig 2D), indicating that the surge in EV secretion was not due to an increase in cell number. Apoptotic bodies derived from dying cells could explain the increasing amounts of particles produced by 3D cultures. However, no signs of apoptosis were detected in the organoids as assessed by cleavage of the apoptosis marker poly(ADP-ribose) polymerase 1 (PARP1) with immunoblotting (Fig 2E).

**Figure 2.**
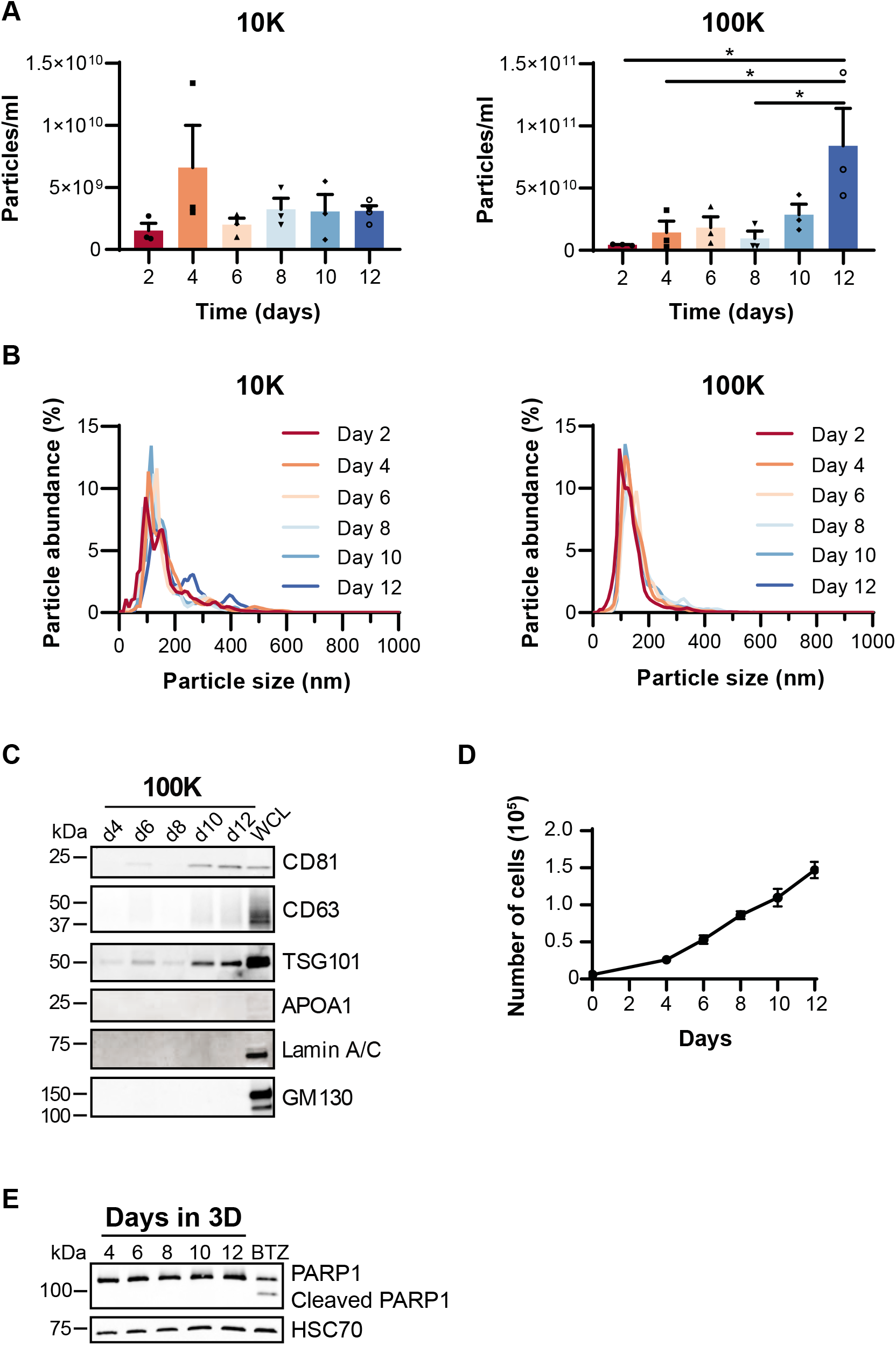
EV secretion increases substantially at the time of invasive transition of PC3 organoids. (A) Particle concentration of the 10K and 100K g pellets obtained by nanoparticle tracking analysis (NTA). Significant differences were assessed by one-way ANOVA with a Tukey post hoc test, *<0.05, +SEM, n=4. (B) EV size distribution of the 10K and 100K pellets detected by NTA shown as mean values of four independent experiments. (C) Immunoblot analysis of EV markers CD81, CD63, and TSG101 in 100K pellets. APOA1, Lamin A/C and GM130 were used as purity controls for the EV samples (d4-d12). WCL, whole cell lysate. (D) Number of PC3 cells grown per well as described in Fig 1. The culture was started (day 0) with 6000 cells per well. Mean values of five independent repeats are shown +SEM. (E) Immunoblot analysis of PARP1 in PC3 cells grown as in Fig 1. Bortezomib (BTZ) treatment (300 nM, 22 h) of 2D grown PC3 cells was used as a positive control for PARP1 cleavage. HSC70 was used as a loading control.

Because Matrigel is a mixture of ECM proteins, it prompted us to examine whether Matrigel-derived particles could interfere with EV analyses. For this purpose, we excluded cells from our experimental setup (Fig 1), and isolated particles from the culture media and dissolved Matrigel. The NTA results demonstrated that although Matrigel released particles of similar size as the cultures with PC3 cells (Supplementary Fig 1D and 3A-C), the amount was only 2-3% of the particles originating from cultures with cells (Supplementary Fig 3D). Importantly, these Matrigel particles did not contain EVs, as assessed by immunoblotting with EV markers TSG101, CD81 and CD63 (Supplementary Fig 3E). Instead, laminin subunit alpha-1 (LAMA1) and 2 (LAMA2), which are two major components of Matrigel (Hughes, Postovit and Lajoie, 2010), were detected. Our findings show that particles originating from Matrigel do not interfere with EV isolation from ECM-based 3D cultures.

The ECM-based 3D culture method described in this study enabled the isolation of EVs from developing organoid cultures. For the first time, EV secretion could be tracked over the course of cancer progression, revealing a surge in the release of EVs, which coincides with the invasive transition of PC3 cells grown in 3D. The increase in EV amounts found at the time of invasion might be essential to tumor progression as EVs have been shown to be able to modify the tumor microenvironment and have a role in pre-metastatic niche formation in metastasis (Zhang and Yu, 2019).

### Proteomic profiling revealed previously undefined EV cargo secreted by PC3 organoids

To investigate if the invasive development of tumors causes qualitative differences in the secreted EVs, we analyzed the size, morphology and protein content of EVs secreted before and after invasive transition. We applied NTA, transmission electron microscopy (TEM), and liquid chromatography-tandem mass spectrometry (LC-MS/MS) analyses for EV characterization. To avoid contaminants in EV samples, such as soluble proteins secreted by the cells or components of the culture media, we used high-resolution iodixanol gradient purification method. The EV isolation with this method depends on mass density of the particles, which results in improved purity of EV samples in comparison to ultracentrifugation (Coumans *et al*., 2017). The EVs were isolated from conditioned media of non-invasive (3D day 2-8), and invasive (3D day 10-14) cultures (Fig 3A). The cultures were expanded to day 14 to gain enough material, whereafter the Matrigel was dissolved to assess if EVs had been trapped therein. For comparison, EVs secreted by 2D PC3 cultures were included in the analyses. From the density gradient purification, 11 fractions were recovered, analyzed by immunoblotting with EV markers (Fig 3B), and the EV-containing fractions 5-7 were pooled together for subsequent analyses. The NTA analyses showed that particle size distribution was similar in all samples, with particle mean sizes ranging between 140 and 169 nm (Fig 4A, Supplementary Fig 4A). This size distribution is in line with the samples isolated with differential centrifugation method (Fig 2B). Furthermore, typical cup-shaped EVs were detected by TEM (Fig 4B), indicating that regardless of the culture conditions, PC3 cells secrete EVs with no differences in size or shape.

**Figure 3.**
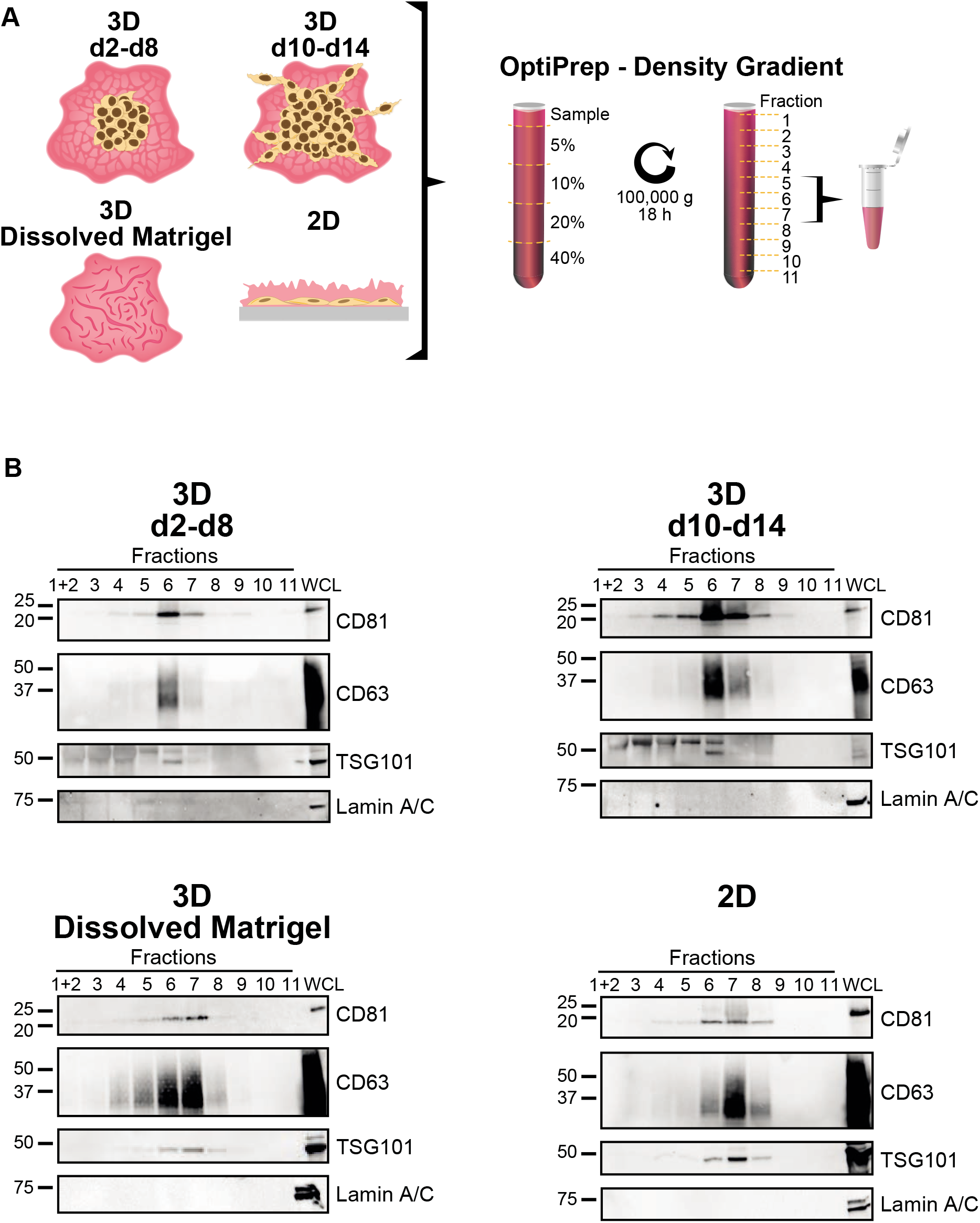
High-resolution gradient purification of EVs from non-invasive and invasive PC3 organoid cultures. (A) EVs were isolated from the conditioned media of non-invasive PC3 3D cultures at days 2-8 (3D d2-d8), invasive cultures at days 10-14 (3D d10-d14), dissolved Matrigel at day 14 (3D Dissolved Matrigel) and conditioned media of 2D cultures (2D). OptiPrep density gradient centrifugation was employed for these samples and the resulting 11 fractions were screened for EVs (B). Fractions 5, 6, and 7 were pooled together and used for subsequent assays (Fig 4–5). (B) Immunoblot analyses of the 11 fractions with EV markers CD81, CD63, and TSG101. Lamin A/C was used as an EV purity marker and PC3 whole cell lysate (WCL) as a positive control for immunoblotting.

**Figure 4.**
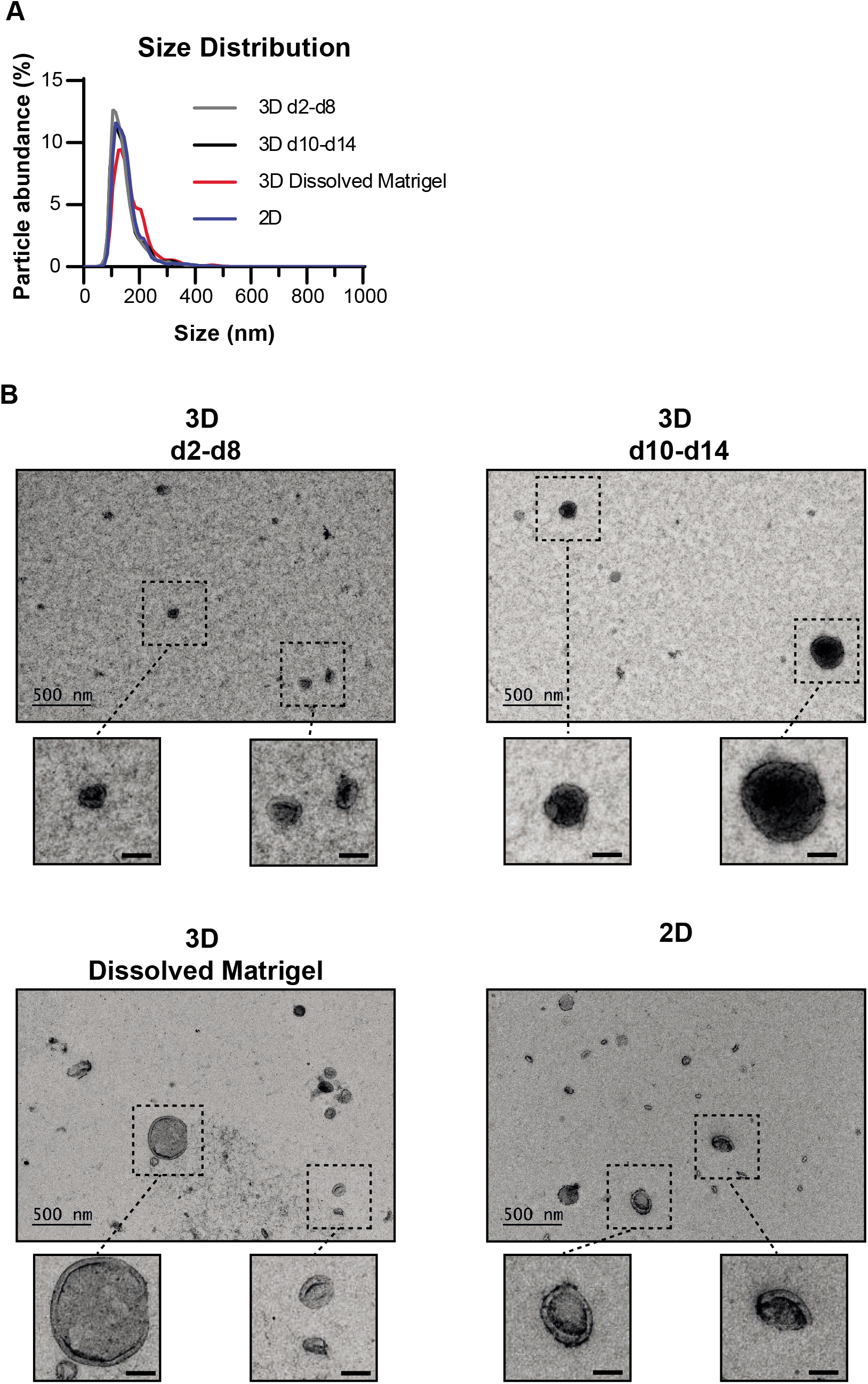
Non-invasive and invasive PC3 organoid cultures secrete EVs of similar size and morphology. (A) **S**ize distribution of EVs from 2D cultures (2D), non-invasive (3D d2-d8) and invasive (3D d10-d14) 3D cultures, and from dissolved Matrigel (3D Dissolved Matrigel). Mean values of three independent experiments conducted with NTA are shown. (B) Morphology of the EVs from 2D and 3D cultures as analyzed by TEM. Representative images (n=2) with scale bar 500 nm for the full sized and 100 nm for the zoomed in images.

The LC-MS/MS analyses revealed extensive differences in protein cargo of EVs secreted by either 2D or 3D cultures (Fig 5A, Supplementary Table 1). This finding is in agreement with previous studies comparing protein content of EVs secreted by 2D and 3D grown cancer cells (Eguchi *et al*., 2018; Rocha *et al*., 2019). Among the 124 identified EV proteins, 56 were unique for 3D cultures, whereas 2D had no unique ones (Fig 5A). Importantly, we identified 27 proteins that have not been previously registered in the Vesiclepedia database for EVs derived from PC3 cells (Kalra *et al*., 2012) (Fig 5B), even though the database contains a comprehensive collection of EV proteins. Nevertheless, the majority of the PC3 EV proteins deposited in Vesiclepedia have been isolated using ultracentrifugation, which might have led to reporting of non-EV proteins, in addition to EV proteins. In conclusion, our data demonstrates that the culture conditions have a profound impact on EV composition, thereby emphasizing the importance of *in vivo*-mimicking models to be used in EV research.

**Figure 5.**
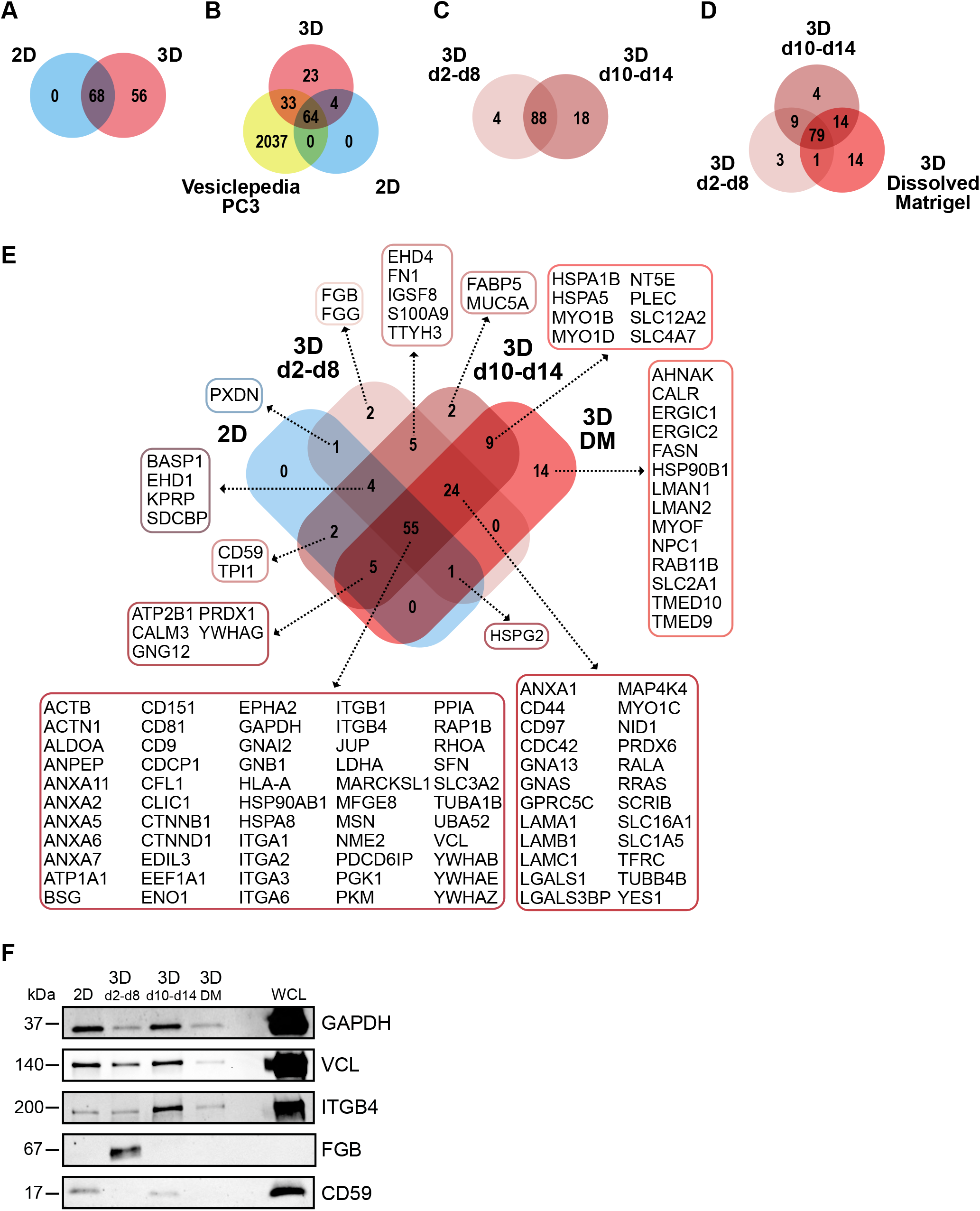
EV protein content changes upon invasive transition of 3D cultured PC3 cells. LC-MS/MS analyses of EV proteins from non-invasive PC3 3D cultures at days 2-8 (3D d2-d8), invasive cultures at days 10-14 (3D d10-d14), dissolved Matrigel at day 14 (3D Dissolved Matrigel) and 2D cultures (2D). (A-D) Venn diagrams representing the number of unique and overlapping EV proteins from PC3 2D and 3D cultures. (B) Comparison of EV proteins from 2D and 3D cultures with EV proteins deposited in the Vesiclepedia database for PC3 cells. (C) Comparison of EV proteins from non-invasive and invasive 3D cultures, (D) and EV proteins retained by the Matrigel (3D Dissolved Matrigel). (E) Venn diagram of the identified EV proteins. (F) Immunoblot analysis of the EV samples with selected MS identified proteins; GAPDH, VCL, ITGB4, FGB and CD59. DM, dissolved Matrigel; WCL, whole cell lysate of 3D grown cells.

### The invasive transition leads to major differences in EV protein composition

Our proteomic profiling strikingly also showed that EV protein content changed upon invasive transition of the PC3 organoids. The EVs secreted by the non-invasive (3D d2-d8) and invasive (3D d10-d14) organoids had distinct protein contents, which became more diversified upon invasion (Fig 5C). Of the identified 124 EV proteins secreted by the 3D cultures, four were unique for the non-invasive and 18 for the invasive cultures. The EVs bound by the Matrigel (3D Dissolved Matrigel) contained 14 unique proteins (Fig 5D). Looking closer at the identified proteins, it became clear that the cancer cells secrete different types of EV cargo proteins before and after invasion (Fig 5E). EVs secreted by invasive organoids (3D d10-d14) included proteins promoting tumor growth (solute carrier family 12 member 2, SLC12A2; sodium bicarbonate cotransporter 3, SLC4A7) (Lee *et al*., 2016; Demian *et al*., 2019), tumor invasion (fatty acid binding protein 5, FABP5) (O’Sullivan and Kaczocha, 2020), and immunosuppression (5’-nucleotidase, NT5E) (Gao, Dong and Zhang, 2014). Interestingly, these proteins were absent from EVs secreted by non-invasive organoids (3D d2-d8). This shows that EVs secreted by invasive organoids contain specific oncogenic signals that can promote several aspects of metastasis. In addition to the distinct EV proteins, all samples derived from 2D or 3D culture conditions contained a common set of classical EV proteins (Théry *et al*., 2018), consisting of tetraspanins, integrins, annexins and heat shock proteins among others (Fig 5E).

To verify the LC-MS/MS results, a subset of the identified proteins was selected for immunoblot analysis (Fig 5F). The results conformed to the expression patterns of the proteins that were analyzed by both methods. Glyceraldehyde-3-phosphate dehydrogenase (GAPDH), vinculin (VCL), and integrin β4 (ITGB4) were detected in all EV samples, which is consistent with the Vesiclepedia database. When comparing the non-invasive and invasive EVs secreted by 3D cultures, we intriguingly found that fibrinogen beta chain (FGB) was only detected in the 3D d2-d8 EV samples. These results show, for the first time, that FGB exists in PC3 EVs. Extracellular FGB is deposited around tumors to function as a scaffold for growth factors and it promotes angiogenesis and tumor growth (Simpson-Haidaris and Rybarczyk, 2001). Since high plasma levels of fibrinogen have been associated with poor cancer prognoses (Zhang and Long, 2017), fluctuations in EV-FGB could serve as an indicator for early stages of cancer. In contrast, CD59 glycoprotein (CD59) was detected only in the 3D d10-d14 samples. Similarly to our results, CD59 has previously been detected in blood plasma (Yan *et al*., 2019) and urine EVs (Lu *et al*., 2009) from prostate cancer patients, which would suggest that CD59 is predominantly a late-stage cargo of prostate cancer EVs.

In summary, we developed an EV production method using ECM-based 3D cultures, which allows tracking of EVs over the course of cancer progression. Using this method, we show that EVs secreted from PC3 prostate cancer cells grown in 2D and 3D cultures have strikingly different protein cargo compositions. Most importantly, the PC3 organoids secrete EVs with previously undefined protein cargo, which changes upon invasive transition. Moreover, the amounts of secreted EVs increased substantially at the time of invasion. These results demonstrate that the culture conditions and the developmental status of the organoid cultures have a major impact on EV secretion and cargo loading, highlighting the necessity of *in vivo*-mimicking conditions for discovery of novel cancer-derived EV components. Our findings set the stage for improved discovery of EV cargo, which can be linked to cancer progression and used as biomarkers for different stages of cancer.

## Methods

### MISEV2018 statement

This work was performed following the minimal information for studies of extracellular vesicles 2018 (MISEV2018) guidelines (Théry *et al*., 2018). We have submitted all relevant data of our experiments to the EV-TRACK knowledgebase (EV-TRACK ID: EV200156) (Van Deun *et al*., 2017).

### 3D cell culture

To collect EVs secreted by PC3 (CRL-1435, ATCC) 3D cultures, we utilized a Matrigel (354230, Corning) sandwich 3D culture system. In brief, an acid-treated (HNO_3_) sterile 13 mm coverslip was placed into every well of a 24-well plate (662160, Greiner). This enabled an even distribution of Matrigel and made the cultures portable and accessible for downstream imaging assays. The plate was chilled and kept cool on a cooling pad during the Matrigel application to ensure even polymerization across the plate. For Matrigel concentration optimization, Matrigel was diluted (3.5, 5.0, and 6.5 mg/ml) in ice-cold serum-free RPMI media (R5886, Merck, with 100 U/μg/ml penicillin/streptomycin [P/S, P0781, Merck] and 2 mM L-glutamine [L-glut, X0550, Biowest]). The Matrigel solutions (50 μl) were drop-casted on the coverslips and the plates were incubated at 37 °C, 5 % CO_2_ for 1 h, during which Matrigel polymerized. PC3 cells (p9-p16) were trypsinized and 6000 cells, counted with Countess II cell counter (Thermo Fisher Scientific), were suspended in 30 μl of EV-free RPMI media (10 % exosome-depleted fetal bovine serum [FBS, A2720801, Thermo Fisher Scientific], 100 U/μg/ml P/S, and 2 mM L-glut) and seeded on each coverslip on top of the Matrigel. The cells were allowed to attach to Matrigel for 1 h in an incubator (37 °C, 5 % CO_2_), whereafter the excess media was removed. A top layer of 30 μl Matrigel (3.5, 5.0, and 6.5 mg/ml) was applied and it was allowed to polymerize for 1 h in the incubator. The wells were then filled with 600 μl of EV-free RPMI media and the plates were placed back in the incubator.

### Media harvest

The conditioned media was carefully harvested every two days (48 h) and replaced with fresh EV-free RPMI media. The media was collected in 15 (188271, Greiner) or 50 ml tubes (210261, Greiner) and was centrifuged twice at 300 g for 10 min at 4 °C in a Sigma 4-16KS centrifuge (Sigma Laboratory Centrifuges) to remove cells and the supernatant was stored at −80 °C until use.

### EV harvest from Dissolved Matrigel (DM)

To collect EVs trapped inside of the Matrigel, the matrix was dissolved at the end of the 3D culture, and the cells were removed. To achieve this, the Matrigel layers were scraped with a scalpel into 8 ml of Gentle Cell Dissociation Reagent (GCDR, 07174, Stemcell) and incubated for 15 min at 37 °C, after which the suspension was mixed by pipetting and placed on a revolver for 15 min at room temperature (RT). To separate the cells from the dissolved Matrigel, the suspensions were centrifuged at 300 g for 10 min at RT. The clear supernatant was collected and the remaining cell-Matrigel pellet was dissolved in 14 ml of phosphate-buffered saline (PBS, L0615, Biowest) and centrifuged at 300 g for 10 min. The supernatants were pooled together as the DM sample. To remove any remaining cells from the sample, the solution was centrifuged twice at 300 g for 10 min at 4 °C. The supernatant and cell pellets were stored at −80 °C until used for EV isolation and cell viability assessments, respectively.

### 2D cell culture

PC3 cells (p9-p16) were plated onto two 15 cm plates in RPMI media (10 % FBS [S181B, Biowest], 100 U/μg/ml P/S and 2 mM L-glut) and were grown in 37 °C, 5 % CO_2_ until they reached 60 % confluency. The cells were then washed with PBS and 15 ml EV-free RPMI (10 % exosome-depleted FBS, 100 U/μg/ml P/S, and 2 mM L-glut) was added. After 48 h cell viability was confirmed by Trypan blue staining (95 %<) and the conditioned media was collected and processed as the 3D cultures.

### EV isolation by differential centrifugation

The EV samples were isolated by differential centrifugation as previously described (Kowal *et al*., 2016). In brief, the samples were thawed at RT and transferred to centrifuge tubes (357003, Beckman Coulter) and centrifuged at 10 000 g in an Avanti J-26 XPI centrifuge (Beckman Coulter) using a JA-25,50 rotor (k-factor 2143.7) for 40 min at 4 °C. The supernatant was transferred to new centrifuge tubes (331372 or 326823, Beckman Coulter) and centrifuged for 90 min at 4 °C in an Optima L-90K ultracentrifuge (Beckman Coulter) at 28 000 rpm (~97 000 g) using an SW41 Ti rotor (k-factor 266). For larger volumes, an SW 32 Ti rotor was used at 29 000 rpm (~103 000 g, k-factor 248). The pellet from 10 000 g centrifugation (10K) was washed with PBS and centrifuged again at 10 000 g for 40 min at 4 °C. The supernatants were discarded and the pellets (10K samples) were dissolved in 100 μl PBS and frozen at −80 °C. The 100K samples’ supernatant was discarded and the pellets were washed with PBS and centrifuged again at 100 000 g for 90 min at 4 °C. The supernatant was discarded and the pellets (100K samples) were re-suspended in 100 μl PBS and frozen at −80 °C.

### Nanoparticle tracking analysis (NTA) of the EV samples

NTA was performed at the EV Core Facility at University of Helsinki using a NanoSight LM14C (Malvern Panalytical) equipped with blue (404 nm, 70 mW) laser and SCMOS camera to characterize particles between 10-1000 nm in the EV samples. The samples were diluted in 0.1 μm filtered (Millex VV, Millipore) PBS to obtain 40-100 particles/view, and five 30 s videos were recorded using camera level 14. The data was analyzed using NTA software 3.0 with the detection threshold 3-4 and screen gain at 10. All NTA experiments were repeated with at least three biological replicates.

### Immunoblotting

Equal volumes of the EV samples were lysed in 5 x RIPA lysis buffer (NaCl 150 mM, Triton X-100 1 %, sodium deoxycholate 0.5 %, SDS 0.1 %, Tris-HCl 50 mM [pH 8.0]) on ice for 1 h and then supplemented with 3x Laemmli sample buffer with or without 2-mercaptoethanol according to antibody preferences. Samples were boiled for 5 min and resolved by SDS-PAGE on 4-20 % Bio-Rad (561094) or Nippon Genetics (PG-S420) gradient gels. After transferring onto a nitrocellulose membrane, the membranes were blocked with 5 % fat-free milk dissolved in 0.05 % PBST for 1 h. The membranes were rinsed with MQ and incubated overnight with primary antibodies (diluted in either PBS or TBS with 0.5 % BSA and 0.02 % NaN_3_) at 4 °C, followed by three PBST or TBST 0.3 % washes and incubation in HRP-conjugated secondary antibodies in blocking solution at room temperature for 1 h. After additional three washes, the signals were detected with enhanced chemiluminescence (34579 and 34094, Thermo Fisher Scientific) and an iBright FL1000 imager (Thermo Fisher Scientific). All immunoblotting experiments were repeated at least three times.

### Antibodies

#### Primary antibodies

Anti-TSG101 (ab125011, 1:1000), anti-GM130 (ab52649, 1:1000), anti-LAMA 1&2 (ab7463, 1:1000), anti-CD59 (ab133707, 1:1000), and anti-GAPDH (ab9485, 1:2500) were purchased from Abcam.

Anti-CD81 (MA5-13548, 1:500) was bought from ThermoFisher Scientific.

Anti-CD63 (CBL553, 1:1000) was acquired from Merck.

Anti-PARP1 (sc-8007, 1:500) anti-FGB (sc-271017, 1:500), anti-APOA1 (sc-376818, 1:100), anti-ITGB4 (sc-514426, 1:300) and anti-VCL (sc-73614, 1:500) was bought from Santa Cruz Biotechnology.

Anti-LMNA (4777, 1:1000) was obtained from Cell Signaling Technology.

Anti-HSC70 (ADI-SPA-815, 1:1000) was purchased from Enzo Life Sciences.

#### Secondary antibodies

Anti-Rabbit IgG (H+L), HRP Conjugate (W4011, 1:5000), and anti-Mouse IgG (H+L), HRP Conjugate (W4021, 1:5000) were purchased from Promega.

Anti-Mouse IgG2a heavy chain (HRP) (ab97245, 1:5000), anti-Mouse IgG2b heavy chain (HRP) (ab97250, 1;5000), and anti-Rat IgG2a H&L (HRP) (ab106783, 1:5000) was bought from Abcam.

### EV isolation by OPTI-prep

In order to obtain enough material for the subsequent assays, the 3D culture was extended to 14 days, thus pooling conditioned media from days 2-8 and days 10-14. The collected media and DM samples were concentrated using a Centricon Plus-70 10K filter (UFC701008, Merck) and diluted to 1 ml using PBS. The concentrates were then layered on top of a high-resolution iodixanol density gradient as previously described (Van Deun *et al*., 2014). In brief, solutions of 5, 10, 20, and 40 % iodixanol were made by mixing appropriate amounts of a homogenization buffer (0.25 M sucrose, 1 mM EDTA, 10 mM Tris-HCL, pH 7.4) and an iodixanol working solution. This working solution was prepared by combining a buffer (0.25 M sucrose, 6 mM EDTA, 60 mM Tris-HCl, pH 7.4) and a stock solution of OptiPrep™ (60 % (w/v) aqueous iodixanol solution, D1556, Merck). The gradient was formed by layering 2.4 ml of 40, 20, 10, and 5 % solutions on top of each other in a 13.2 ml open-top polyallomer tube (331372, Beckman Coulter). The 1 ml concentrates were overlaid on top of the gradient, and centrifuged at 100 000 g (28 000 rpm, k-factor 266) in an Optima L-90K ultracentrifuge (Beckman Coulter) using an SW41 Ti rotor for 18 h at 4 °C with slow deacceleration. The gradient was separated from the top into 1 ml fractions, diluted to 11 ml with PBS, and centrifuged at 100 000 g for 3 h at 4 °C with max deacceleration. The acquired pellets were diluted in 250-500 μl of PBS, split into smaller aliquots for downstream assays, and stored at −80 °C.

### EV lysis, in-solution digestion, and LC-MS/MS

Isolated EVs were lysed in a buffer containing 8 M urea, 0.5 % NP40, 2 mM EDTA, 150 mM NaCl and one tablet of protease and phosphatase inhibitors (A32959, Thermo Fisher Scientific) for 30 min on ice. Then, the samples were sonicated for 5 min (30’ on and 30’ off) and the proteins were precipitated using 4-5 volumes of cold acetone overnight at −20 °C. Samples were cleared by centrifugation for 15 min using 16 000 g at 4 °C and the pellet was dissolved in 100 μl of 6 M urea in 25 mM ammonium bicarbonate. Approximately 100 μg of protein was used for in-solution digestion. The samples were reduced with 10 mM DTT for 1 h at 37 °C and alkylated with 40 mM iodoacetamide for 1 h in the dark. The alkylation was quenched with 40 mM DTT and the urea concentration was diluted by adding 900 μl of 25 mM ammonium bicarbonate. Sequencing grade trypsin was added to each sample in a 1:30 ratio to total protein and the samples were incubated overnight at 37 °C. The peptides were acidified with TFA and were desalted using Sep Pak C18 columns (100 mg, Waters) according to the instructions of the manufacturer. The samples were dried on a speed-vac centrifuge and stored at −80 °C. Before analysis, the samples were dissolved in 0.1 % formic acid and approximately 100 ng of each sample was loaded for LC-MS/MS analysis. The analysis was conducted using an Easy-nLC 1000 liquid chromatograph (Thermo Fisher Scientific) coupled to an Orbitrap Fusion Lumos Tribrid Mass Spectrometer (Thermo Fisher Scientific). The peptides were loaded on a pre-column (100 μm × 2 cm), followed by separation in an analytical column (75 μm x 15 cm), both packed with 5 μm ReproSil-Pur 200 Å C18 silica particles (Dr. Maisch HPLC GmbH). The peptides were separated using a 60 min gradient (5-42 % B in 50min, 42-100 % B in 6 min, 100 % B for 4 min, in which solvent B was 80 % acetonitrile 0.1 % formic acid in water) at a flow rate of 300 nl/min. MS/MS data were acquired in positive ionization mode using data-dependent acquisition using a 2.5 s cycle time. The MS survey scans were acquired with a resolution of 120 000 with the range of 300-1700 m/z, an AGC target of 7.0 e5, and a maximum injection time of 50 ms. The ions were selected for HCD fragmentation using an isolation window of 1.6 m/z, with AGC target of 104 and a maximum injection time of 50 ms. MS/MS spectra were recorded with 30 000 resolution, and a dynamic exclusion window of 35 s was used.

### Proteomic data processing

Raw files obtained from the LC-MS/MS analyses were processed using MaxQuant software (version 1.6.7.0) (Tyanova, Temu and Cox, 2016) and searched against a SwissProt human protein database (https://www.uniprot.org/, release 20/10/2019) with added common contaminants using the built-in Andromeda search engine. Trypsin digestion with a maximum of 2 missed cleavages, cysteine carbamidomethylation as a fixed modification, and methionine oxidation as variable modification were selected as the parameters of these searches. “Match between runs” option in MaxQuant was used with a matching time window of 2 min and an alignment time window of 20 min. The peptide level false discovery rate (FDR) was set to 1 % and was determined by searching against a concatenated normal and reversed sequence database.

Label-free quantitation was performed using the fast LFQ algorithm. Otherwise, the default settings in MaxQuant were used in data processing. The database search results with LFQ intensities were analyzed using Perseus (version 1.6.7.0) (Tyanova *et al*., 2016). Proteins only identified by site, proteins considered contaminants, and reverse were discarded. Proteins identified with <2 unique peptides were filtered out and only proteins with 2 values in at least one group were kept. Missing values were not imputed. The results are represented as log2 (LFQ) of two independent replicates. Venn diagrams were made using Funrich (Pathan *et al*., 2015) (last updated 11.11.2020). Supplementary Table 1 contains all the identifications and transformed LFQ values.

### Transmission electron microscopy (TEM)

TEM sample preparation and imaging were performed by the EV Core Facility, University of Helsinki, essentially as previously described (Puhka *et al*., 2017). In brief, pioloform- and carbon-coated copper grids were glow discharged before the samples were loaded. The grids and samples were fixed with 2 % paraformaldehyde in 0.1 M NaPO4 buffer (pH 7.0), stained with 2 % neutral uranyl acetate, and further stained and embedded in uranyl acetate and methyl cellulose mixture (1.8/0.4 %). The EVs were viewed using Jeol JEM-1400 (Jeol Ltd) operating at 80 kV. Images were acquired with Gatan Orius SC 1000B CCD-camera (Gatan Inc).

### Statistics and reproducibility

All experiments were repeated with at least three biological repeats unless otherwise stated. Data are expressed as mean±SEM. For multiple group comparisons, one-way ANOVA or two-way ANOVA, followed by a Tukey test, was performed. A p-value < 0.05 was considered statistically significant. GraphPad Prism 8 was used for statistical analysis.

### 3D culture cell counting

After harvesting media from 3D cultures, coverslips were collected and broken into 14 ml of ice-cold PBS supplemented with 5 mM of EDTA. The suspensions were mixed by rotation for 45 min at 4 °C to fully dissolve the Matrigel, followed by centrifugation at 200 g for 5 min at 4 °C. The supernatant was removed and the cells were washed with 14 ml cold PBS, mixed, and recentrifuged at 200 g for 5 min at 4 °C. The supernatant was removed and the cell pellets were diluted in 2 ml trypsin (L0931, Biowest) (37 °C) and incubated for 15 min at 37 °C, agitating the tubes every few minutes. After the incubation, 10 ml of RPMI (10 % FBS, 100 U/μg/ml P/S, and 2 mM L-glut) was added to the tubes and the cells were pelleted by 5 min centrifugation at 300 g at RT. The cells were resuspended in 100 μl of media and counted using the Countess II cell counter (Thermo Fisher Scientific). Four wells were counted for each time point to obtain an average cell amount per well. The experiments were repeated five times.

### Brightfield imaging of 3D cultures

The 3D cultures were imaged from multiple wells every 48 h using a ZEISS Axio Vert.A1 microscope with an Axiocam 506 camera and an LD A-Plan 5x/0.15 Ph1 objective. Brightness and contrast were adjusted using ZEN 2012 (ZEISS). Adjustments were applied across the entire images without loss of data.

## Acknowledgements

We gratefully acknowledge Diosángeles Soto Véliz for her graphical design used in Figures 1 and 3A. Malin Åkerfelt and Jenny Joutsen, together with all members of the Sistonen laboratory, are thanked for their valuable comments and critical review of the manuscript. We thank the Proteomics Facility of Turku Bioscience Centre (University of Turku and Åbo Akademi University) for use of equipment and the EV Core Facility (University of Helsinki) for conducting the NTA and TEM analyses and providing expert comments. This project was funded by Åbo Akademi University (JCL, LS, EH), Academy of Finland (LS), Liv och Hälsa (LS, EH), Sigrid Jusélius Foundation (LS), Cancer Society of Finland (LS), Otto A. Malm Foundation (EH), Magnus Ehrnrooth Foundation (LS, EH), Swedish Cultural Foundation (JCL, EH), and K. Albin Johansson Foundation (EH).

## Author Contributions

JCL and EH designed the experiments; JCL, LSCR, SHB, JR, and EH performed the experiments; JCL, LSCR, and EH analyzed the data; JCL, LS, and EH wrote the manuscript with input from all other authors.

## Competing Interests

The authors declare that they have no conflict of interest.

---

**Supplementary Figure 1 - Matrigel concentration affects EV isolation efficiency.** (A) Representative brightfield images of PC3 cells grown as in Fig 1 in Matrigel (3.5, 5.0 or 6.5 mg/ml) for 12 days. Scale bar 200 μm. (B-D) The size and concentration of particles isolated from conditioned media (day 10 to day 12) from PC3 cell cultures by 10K and 100K g centrifugation and analyzed by NTA. The used Matrigel concentrations were 3.5, 5.0, and 6.5 mg/ml. The data are presented as mean values of four independent experiments. (C) Significant differences were assessed by one-way ANOVA and Tukey’s multiple comparison test, *p<0.05, +SEM.

**Supplementary Figure 2 - Particle characteristics of EVs secreted by 3D grown PC3 cells.** PC3 cells were cultured and EVs were isolated as described in Fig 1. The NTA data presents mean and mode values of 10K and 100K pelleted particles of four independent experiments.

**Supplementary Figure 3 - Matrigel-derived particles do not impede EV isolation.** Cultures without PC3 cells in 3.5 mg/ml of Matrigel were done as described in Fig 1. Media was collected at day 12 and dissolved Matrigel at day 14, and particles were isolated by differential centrifugation (10K and 100K g). The obtained results were compared to 3D cultures with cells. (A-C) NTA of media or dissolved Matrigel from conditions without cells. The data are presented as mean values of three independent experiments. (B) Significant differences were assessed by one-way ANOVA with Tukey’s multiple comparison test, ns=not significant, *p<0.05, +SEM, n=4. (D) NTA of media or dissolved Matrigel from cultures with and without cells (days 10 to 12). The data are presented as mean values of three independent experiments. Significant differences were assessed by two-way ANOVA with Tukey’ s multiple comparisons test, ns=not significant, *p<0.05, **p<0.005, +SEM, n=4. Pie chart: particle amount comparison from media or dissolved Matrigel with and without cells (gray and black, respectively). The 10K and 100K sample particles were combined in this comparison. (E) Immunoblot analysis of EV markers TSG101, CD81 and CD63, and ECM components LAMA1 and LAMA2 in media and dissolved Matrigel (DM) from conditions without cells. EVs from conditioned media (EVs) and whole cell lysate (WCL) of 2D cultured PC3 cells were used as positive controls.

**Supplementary Figure 4. Non-invasive and invasive PC3 organoid cultures secrete EVs of similar size.** (A) NTA results of EVs from non-invasive PC3 3D cultures at days 2-8 (3D d2-d8), invasive cultures at days 10-14 (3D d10-d14), dissolved Matrigel at day 14 (3D Dissolved Matrigel) and 2D cultures (2D). The data presents mean values of three independent experiments. (B) Minimum and maximum EV sizes (nm, diameter) observed by TEM from the EV samples. The data presents mean values of two independent experiments

